# Optimizing Real Time fMRI Neurofeedback for Therapeutic Discovery and Development

**DOI:** 10.1101/003400

**Authors:** L. E. Stoeckel, K. A. Garrison, S. Ghosh, P. Wighton, C. A. Hanlon, J. M. Gilman, S. Greer, N. B. Turk-Browne, M. T. deBettencourt, D. Scheinost, C. Craddock, T. Thompson, V. Calderon, C. C. Bauer, M. George, H. C. Breiter, S. Whitfield-Gabrieli, J. D. Gabrieli, S.M. LaConte, L. Hirshberg, J. A. Brewer, M. Hampson, A. Van Der Kouwe, S. Mackey, A. E. Evins

**Affiliations:** MGH Department of Psychiatry; Harvard Medical School; Athinoula A. Martinos Center; McGovern Institute for Brain Research (MIT); Yale University School of Medicine, Department of Psychiatry; MGH Department of Radiology; Medical University of South Carolina, Department of Psychiatry; University of California at Berkeley, Department of Neuroscience; Princeton University, Department of Psychology; Princeton Neuroscience Institute; Yale University School of Medicine, Department of Diagnostic Radiology; Child Mind Institute; Universidad Nacional Autonoma de Mexico, Instituto de Neurobiologia; Northwestern University Feinberg School of Medicine, Department of Psychiatry and Behavioral Sciences; Virginia Tech, School of Biomedical Engineering and Sciences; Virginia Tech Carilion Research Institute; Alpert Medical School of Brown University, Department of Psychiatry and Human Behavior; University of Massachusetts Medical School, Department of Medicine and Psychiatry; Stanford University School of Medicine, Department of Anesthesia

## Abstract

While reducing the burden of brain disorders remains a top priority of organizations like the World Health Organization and National Institutes of Health (BRAIN, 2013), the development of novel, safe and effective treatments for brain disorders has been slow. In this paper, we describe the state of the science for an emerging technology, real time functional magnetic resonance imaging (rtfMRI) neurofeedback, in clinical neurotherapeutics. We review the scientific potential of rtfMRI and outline research strategies to optimize the development and application of rtfMRI neurofeedback as a next generation therapeutic tool. We propose that rtfMRI can be used to address a broad range of clinical problems by improving our understanding of brain-behavior relationships in order to develop more specific and effective interventions for individuals with brain disorders. We focus on the use of rtfMRI neurofeedback as a clinical neurotherapeutic tool to drive plasticity in brain function, cognition, and behavior. Our overall goal is for rtfMRI to advance personalized assessment and intervention approaches to enhance resilience and reduce morbidity by correcting maladaptive patterns of brain function in those with brain disorders.

## Introduction

Researchers have recently developed neuroimaging technologies that provide us with powerful tools to better understand the complexity of human brain-behavior relationships with the goal of discovering and developing novel, safe, effective and personalized therapeutics to treat brain disorders. Recognizing the potential of these new tools for advancing clinical neuroscience, the European Union and United States launched the Human Brain Project and Brain Research through Advancing Innovative Neurotechnologies (BRAIN) initiatives with estimated budgets of $1.3 billion and $4.5 billion in research support, respectively, to accelerate the development of such neurotechnologies (BRAIN, 2013; Kandel et al., 2013). At the leading edge of neuroimaging technology development is real time functional magnetic resonance imaging (rtfMRI), which allows a non-invasive view of brain function^1^ *in vivo* and in *real time*^2^. rtfMRI has the potential to be used as a clinical neuroimaging tool in diagnosis, monitoring of disease, tracking of therapeutic response, and uniquely, in treatment itself via rtfMRI neurofeedback. rtfMRI neurofeedback is an application of this technology that can be used to assess and/or alter patterns of brain activity associated with cognition or behavior while an individual is inside the MRI scanner in real time (Birbaumer et al., 2009; Birbaumer et al., 2006; deCharms, 2008; deCharms et al., 2004; deCharms et al., 2005; Weiskopf et al., 2007; Weiskopf et al., 2003). The therapeutic potential for this approach lies in its ability to alter neural plasticity and learned behavior to modify brain function to optimize and/or restore healthy cognition and behavior.

Brain structure and function are modified in response to changes within and outside the central nervous system via both normal and disordered processes (Kolb et al., 2010). Compared to standard fMRI experiments in which behavior is manipulated and subsequent changes in brain activity are measured, rtfMRI switches the direction of the relationship between brain and behavior so that we can determine if *directly* changing brain function leads to changes in behavioral or experiential outcomes (Weiskopf, 2012). This approach of facilitating specific changes in brain function to produce changes in cognition, experience, or behavior is theorized to occur by skill learning (Birbaumer et al., 2013) and the acceleration and optimization of systems-level neuroplasticity (Sagi et al., 2012), as has been observed with other brain-based training protocols (Anguera et al., 2013). Other neurotherapeutic technologies, including electroconvulsive therapy, vagus nerve stimulation, deep brain stimulation, and transcranial magnetic stimulation or transcranial direct current stimulation, are also being used or investigated for the treatment of brain disorders and may also produce clinical change via altered neuroplasticity. Each of these technologies has potential benefits and may also be limited by constraints in spatial resolution or by their invasive nature.

Neurofeedback is a training method in which real time information about changes in neural activity is provided to an individual to facilitate learned self-regulation of this neural activity to produce changes in brain function, cognition, or behavior. The earliest studies of neurofeedback employed electroencephalography (EEG) and demonstrated feedback-related changes in electrical brain activity and related behavior and cognition in humans (Keizer et al., 2010; Kouijzer et al., 2009; Ros et al., 2013; Zoefel et al., 2011) and other animals (Philippens and Vanwersch, 2010; Schafer and Moore, 2011; Sterman et al., 1969). Brain change after EEG neurofeedback has been shown using EEG and event related potentials (Egner and Gruzelier, 2001; Kropotov et al., 2005). Likewise, changes in fMRI response after EEG neurofeedback have been shown in targeted neural networks after a single 30-minute EEG training session (Ros et al., 2013) and in specific symptom-related brain regions of interest (ROI) after multiple training sessions (Levesque et al., 2006). There have been several randomized controlled trials (RCTs) using EEG-based feedback, primarily in patients with attention deficit hyperactivity disorder (ADHD) (Hirshberg et al., 2005). A recent meta-analysis of existing RCTs indicates that EEG feedback training is associated with a reduction of ADHD symptoms with a large effect size (Arns et al., 2009) and a large randomized, sham-controlled trial is currently underway (LH, personal communication). However, while EEG has superior temporal resolution compared to standard fMRI, poor spatial resolution including the so-called ‘inverse problem’ (Kriegeskorte et al., 2009) limit the clinical utility of EEG. By contrast, rtfMRI can be used to target brain regions and networks with improved anatomical precision beyond EEG and improved temporal resolution beyond standard block design fMRI. Finally, rtfMRI and EEG neurofeedback can be used simultaneously to take advantage of the spatial resolution of fMRI and the temporal resolution of EEG with the hope that this combined approach will lead to more efficient neuroadaptive changes and more effective clinical outcomes (Zotev et al., 2014).

rtfMRI was developed in 1995 (Cox et al., 1995), and proof-of-concept for rtfMRI as a potential neurotherapeutic tool for the treatment of brain disorders was demonstrated in 2005 (deCharms et al., 2005). There have since been substantial advancements related to rtfMRI technology and implementation (Hinds et al., 2011; LaConte, 2011; Weiskopf et al., 2005), with reports of rtfMRI modification of function in several brain structures. Although rtfMRI has multiple potential applications as a clinical neuroimaging tool, the research to date has been focused on the use of rtfMRI neurofeedback to alter brain function and behavior. From this research, several groups have reported successful application of rtfMRI to modify cognitive and behavioral processes relevant for the treatment of clinical disorders (for review of these studies see Birbaumer et al., 2009; Caria et al., 2012; Chapin et al., 2012; deCharms, 2007; deCharms, 2008; Sulzer et al., 2013a; Weiskopf, 2012; Weiskopf et al., 2007). Studies have demonstrated promise of rtfMRI neurofeedback in the treatment of chronic pain (deCharms et al., 2005), tinnitus (Haller et al., 2010), stroke (Sitaram et al., 2012), depression (Linden et al., 2012), schizophrenia (Ruiz et al., 2013), obesity (Frank et al., 2012), and addiction (Hartwell et al., 2013; Li et al., 2013). Given the early stage of this research, it is not surprising that there are many limitations to these studies. Most notably, small sample sizes (typically between 6-12 participants) and lack of critical control conditions limit their potential use as evidence-based interventions. There are several plausible alternative hypotheses for the key variable(s) that account for the changes observed following rtfMRI neurofeedback training. These include, but are not limited to, effects due to experimenter monitoring, self-monitoring, positive reinforcement, cognitive and emotion regulation strategies, enhanced self-efficacy and motivation to change due to successful performance, and placebo response. To date, there have been no large RCTs of rtfMRI neurofeedback. RCTs involve random allocation of participants to treatment and control groups, minimizing bias in treatment assignment and facilitating concealment of treatment assignment to experimenters and participants (Schulz and Grimes, 2002). RCTs are the gold standard for ‘rational therapeutics’ in clinical medicine (Meldrum, 2000) and are critical for establishing an evidence-based clinical practice (The Cochrane Collaboration, 2011). Nevertheless, this important early work supports the investment in RCTs of rtfMRI for the treatment of some brain disorders.

The aim of this paper is to define guidelines to establish the *therapeutic* utility of rtfMRI neurofeedback by emphasizing clinical issues that should be considered beyond the technical considerations that have been the primary focus of more recent reviews (Birbaumer et al., 2013; Sulzer et al., 2013a). Although the guidelines were developed with regard to the use of rtfMRI neurofeedback to treat brain disorders, resolution of these issues is also necessary to advance development of rtfMRI as a tool in clinical neuroimaging more generally. For each guideline, we delineate the challenges and potential limitations of rtfMRI that need to be addressed to advance development of this neurotechnology, and outline a research strategy to address these challenges and limitations including potential experimental and neuroinformatics approaches.

## Guidelines for establishing real time fMRI as a neurotherapeutic tool

Several technical, neuroscientific, and clinical issues must be addressed before rtfMRI neurofeedback can advance as a clinical neuroimaging tool, and in particular for direct therapeutic applications. Due to the complexity of rtfMRI neurofeedback experiments, it is advisable to have considered these issues and have solutions in place in order to maximize the likelihood that the experiment will be a success. We have outlined the issues we felt are most critical to clinical applications of rtfMRI neurofeedback and have offered potential solutions to help guide researchers.

### Guideline 1: The rtfMRI signal is accurate and reliable

Necessary preconditions for any successful rtfMRI experiment are that the brain state of an individual is detectable and can be reliably and reproducibly converted into a feedback signal over the time-scale in question. Here we propose possible metrics that can be used to evaluate these prerequisites.

From a methodological perspective, the neurofeedback signal is generally derived from fMRI paradigms of two broad categories: general linear model (GLM)-based methods, and more recently, multivariate pattern analysis (MVPA) methods. For recent reviews of these methods, see Sulzer (Sulzer et al., 2013a) and LaConte (LaConte, 2011), respectively. GLM-based methods define an *a priori* ROI, either anatomically (using anatomical landmarks or atlas-based techniques) or with a functional localizer. The GLM is used to regress out nuisance parameters, and the resulting BOLD signal at each voxel in the ROI is combined into a neurofeedback signal using either averaging or a weighted average based on the standard deviation of the residual of the GLM in each voxel (Hinds et al., 2011). MVPA methods use supervised learning techniques, usually support vector machines, to determine the optimal set of weights (from either the whole brain or a restricted ROI) used to combine the BOLD signal across voxels into a single neurofeedback score.

In the context of rtfMRI, ‘real time’ is often used to describe both neurofeedback signals estimated from a single brain volume acquisition (Hinds et al., 2011) and across several brain volume acquisitions ((Bohlmeijer et al., 2011; Cox et al., 1995; Johnson et al., 2012; Yoo et al., 1999) to mitigate the contribution of noise in the BOLD signal. In order to estimate a reliable BOLD signal in a single measurement, it is important for models to include a moment-to-moment estimate of noise. One example of this approach is the incremental GLM method described in (Hinds et al., 2011) and implemented in murfi2, software that is freely available on Github (http://github.com/gablab/murfi2).

Regardless of whether a GLM- or MVPA-based model is used to compute the neurofeedback signal or whether the signal is estimated in a single brain volume acquisition or across several acquisitions, the signal is a one-dimensional, usually linear combination of the BOLD signal across the brain. We can use the signal to noise ratio (SNR) to compute how well the neurofeedback signal conforms to the experimental design.

Let F represent the (stochastic) neurofeedback signal and X the (deterministic) experimental design vector. Under the standard GLM assumptions (Monti, 2011) SNR can be calculated by:

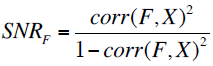

Where corr(F,X) represents the Pearson correlation coefficient of vectors F and X. This quantity can be used to estimate how many repetition times (TRs) the neurofeedback signal would need to be averaged over to ensure it is accurate with a confidence level of (1−*α*_*F*_).

After theoretical modeling, we arrived at an equation that relates three key quantities: the confidence in the neurofeedback signal (1−*α*_*F*_), the signal to noise ratio (*SNR*_*F*_), and the number of TRs used to compute the neurofeedback signal (n). The relationship between these 3 variables is:

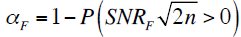

When P() is the standard cumulative normal distribution and n is the length (in TRs) of both the task and fixation blocks.

Using this formula, we can determine the block length necessary to estimate an accurate neurofeedback signal with 95% confidence. We tested this formula using three rtfMRI feedback paradigms with regional ROIs with small (ventral striatum or VS), medium (fusiform face area and parahippocampal place area or FFA/PPA), and large (somatomotor cortex or SMC) expected SNR values. The results are illustrated in Figure 1 and summarized in Table 1.

**Figure 1.**
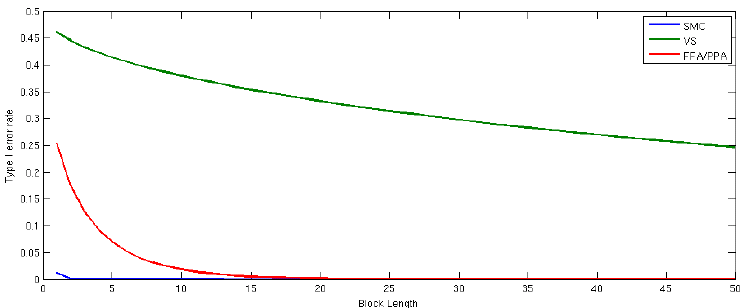
The relationship between signal-to-noise ratio (SNR) and block length (n) required for a type I error rate (α) less than 0.05 for three ROIs.

**Table 1.** The relationship between signal-to-noise ratio (SNR) and block length (n) required for a type I error rate (α) less than 0.05 for three ROIs.

| ROI | SNR | n required for $\alpha < 0.05$ |
| --- | --- | --- |
| SMC | 1.581 | 1 |
| VS | 0.0685 | 289 |
| FFA/PPA | 0.4668 | 7 |

Equally important to a detectable signal is a reproducible signal. Here the concordance correlation coefficient can be used to determine the reproducibility of the rtfMRI signal for a given subject from one run to the next, given the model employed. The concordance correlation coefficient is a simple metric that has been applied to fMRI to evaluate repeatability of various models (Lange et al., 1999). Let W_1_ and W_2_ represent the weights used to aggregate the feedback signal (derived from either a GLM or MVPA-based model) from runs 1 and 2, respectively. The concordance correlation coefficient between these weight vectors is then

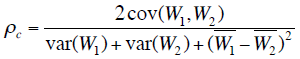

Where cov(.) represents the covariance of the weight vectors, var(.) represents the variance and 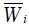 represents the mean.

From this work, we determined that the FFA/PPA and SMC ROIs were feasible as neurofeedback brain regions to target, but VS with low SNR was not feasible. It should be emphasized that detection of activation in FFA/PPA and SMC ROIs is feasible *relative* to the VS ROI; however, there are still substantial challenges in estimating neural activity from a single noisy BOLD volume (Hinds et al., 2011) that should be considered when deciding whether moment-to-moment neurofeedback from an ROI, set of ROIs, or components will produce the intended study outcome. At a minimum, it is recommended that researchers establish the quality of the neurofeedback signal in an independent dataset using either the SNR and concordance coefficient approach described above or some analogous method. If an experimenter finds that a signal does not meet these minimum standards, efforts should be made to optimize the experimental parameters and/or target ROI(s) before collecting data for a larger planned study. In cases where neurofeedback occurs from a region or network with low SNR, e.g., (Sulzer et al., 2013b), reporting SNR and methods used to optimize SNR, will help guide future researchers targeting this brain areas for neurofeedback.

### Guideline 2: rtfMRI neurofeedback leads to learning

One use of rtfMRI is to provide feedback aimed at inducing learning that is difficult to achieve, or is less efficient, using other methods. This can be contrasted with alternative uses of rtfMRI, such as triggering task events or optimizing task parameters, where the goal is not necessarily to produce a lasting learning effect. To understand the learning induced by rtfMRI neurofeedback, three aspects of this process should be considered: (1) How is learning measured? (2) Which mechanisms are responsible? and (3) How meaningful or lasting are the effects? We will describe these three aspects followed by a case study.

Learning can be said to occur when experience influences behavior and/or alters brain structure or function. In the case of rtfMRI neurofeedback, the relevant experience can consist of elements of the task, feedback about regional brain activation, multivariate patterns of brain activity, or connectivity, and cognitive processes that arise due to the task, feedback, or attempts to control feedback with strategies. The consequences of this experience can be assessed using various dependent measures linked to learning in more standard experiments in cognitive psychology and neuroscience. In behavior, learning can be reflected in improved perception (Fahle, 2002), memory recall and recognition (Yonelinas, 2002), anticipation/prediction (Bubic et al., 2010), priming (Tulving and Schacter, 1990), and motor action (Stadler and Frensch, 1998). In the brain (particularly fMRI), learning can be reflected in enhanced (Schwartz et al., 2002) or attenuated activation within sensory systems (Grill-Spector et al., 2006; Turk-Browne et al., 2008), activation in learning and memory systems (Brewer et al., 1998; Poldrack et al., 2001; Wagner et al., 1998), changes to the multivariate representational space of brain regions (Folstein et al., 2013; Schapiro et al., 2013), changes to functional connectivity (Buchel et al., 1999), increased gray matter volume (Draganski et al., 2004), and alterations in white matter (Zatorre et al., 2012). All of these measures are potential targets for rtfMRI studies seeking to induce learning with neurofeedback.

These changes in behavior and the brain reflecting learning can arise from different mechanisms. One proposed mechanism is reward-based skill learning via cortico-basal ganglia brain networks (Birbaumer et al., 2013). Feedback from a brain region may produce a type of instrumental conditioning, whereby activation of that region becomes rewarding. An increase in the activity of the region, and possibly additional inputs from the reward system and regions involved in cognitive control, may induce local plasticity. This plasticity could be reflected in alterations of the selectivity and circuitry of neurons in that region (Sur and Rubenstein, 2005). There may also be larger-scale consequences. The region/representation used as the basis for feedback may become more involved in general, or more selective for an ongoing task. This could be analogous to establishing a compensatory mechanism, as occurs naturally after brain damage or in aging (Bedny et al., 2011; Heuninckx et al., 2008). However, the region(s) being “compensated” for (i.e., initially involved but not used for neurofeedback) remain intact, and could possibly be recruited less over time. Other regions that implement cognitive strategies for controlling feedback may become engaged in addition to the region being targeted for neurofeedback and the other brain areas recruited by this target region(s).

These learning effects differ in several ways that will impact the likelihood of obtaining an effect, where the effect will be observed in the brain, and whether the effect will be manifested in behavior. Several characteristics of learning will determine the feasibility of an rtfMRI study. For instance, learning occurs over a range of timescales, from immediately in the case of priming to over weeks in the case of perceptual learning. Relatedly, effects persist for different durations depending on the type of learning and brain system involved, from milliseconds for adaptation in the visual system (Grill-Spector et al., 2006) to years for episodic memories consolidated in association cortex (Tomasino et al., 2012). Finally, learning procedures vary in terms of whether the effects generalize to other contexts from being hyperspecific to the training context (Jiang and Song, 2005) to more flexible (Turk-Browne and Scholl, 2009). Considering these parameters when designing a study will be important for its success.

These three aspects of learning –dependent measures, neural mechanisms, and timing/generalization properties– allow rtfMRI neurofeedback studies to be classified, and opportunities for new research to be identified. Take, for example, a study of perceptual learning induced by rtfMRI (Shibata et al., 2011). The dependent measures in this study were improved (1) accuracy in identifying a trained visual target orientation and (2) multivariate classification of this orientation from early visual cortex. The proposed mechanism is that rtfMRI feedback encouraged participants to generate orientation-specific neural activity patterns in this region, which resulted in local plasticity. The properties of learning in this task are that it took 5-10 sessions of training over a month, the effects lasted at least that long, and the benefit of training was specific to one orientation at a particular contrast. Various aspects of this classification could be investigated in future studies, such as what other brain systems are responsible for creating the activity patterns in visual cortex, and whether other forms of perceptual learning (e.g., Xiao et al., 2008) could lead to more generalized benefits.

### Guideline 3: The training protocol is optimized for rtfMRI-based neurofeedback and learning

A variety of approaches have been used for training subjects to control their brain patterns via rtfMRI neurofeedback. As it is yet unclear which approach yields optimal learning, more work is needed to provide evidence-based guidelines for clinical trial design. In most studies, the nature of the neurofeedback signal is made explicit to the subject, but there have also been paradigms where training occurs covertly (Bray et al., 2007; Shibata et al., 2011). Some studies present feedback while subjects are processing auditory, visual, or tactile stimuli (Bruhl et al., 2014; deCharms et al., 2005; Scheinost et al., 2013; Yoo et al., 2007) or while they are performing an assigned cognitive task (Chiew et al., 2012). Other rtfMRI studies employ an unconstrained paradigm in which the neurofeedback and the cues to increase or decrease brain activity are the only stimuli provided and the subjects are free to use any cognitive strategy to control brain function and neurofeedback (Caria et al., 2007; Hampson et al., 2011; Rota et al., 2009). The optimal approach is likely dependent upon the application.

One open question is the importance of providing initial support (e.g., neurostimulation, pharmacotherapy, computerized cognitive training, and/or cognitive strategies) to subjects that will enable them to exert some initial level of control over the relevant brain activity patterns. Early reports suggested that learning was greatly facilitated by providing cognitive strategies to the subjects before they began neurofeedback (Caria et al., 2007; deCharms et al., 2005). Generally, rtfMRI neurofeedback studies have either provided all subjects with strategies or have not provided any subjects with strategies, making it impossible to determine the degree to which discussing strategies with subjects before they began neurofeedback helped the subjects to gain control over their patterns of brain activity. There have been no published studies to date that have used neurostimulation, pharmacological aids, or computerized cognitive training to enhance subject’s ability to utilize rtfMRI feedback, which may be an important avenue for future studies to explore.

It is not always necessary to provide initial strategies to subjects in order to achieve learning. Successful learning was achieved in subjects who were trained using instrumental conditioning (Bray et al., 2007) or were not provided with any initial strategies (Shibata et al., 2011). Further, subjects who have learned to control their brain activity patterns via rtfMRI neurofeedback have not always been consciously aware of the mental functions that were being molded by the training, even after completion of the neurofeedback (Shibata et al., 2011). These data imply that rtfMRI neurofeedback can induce subconscious learning. Although this is encouraging in terms of the potential utility of rtfMRI for training mental function, it also has ethical implications that must be carefully considered.

It is also important to consider the schedule of delivery of neurofeedback, whether it is continuous or intermittent, in order to optimize learning. Continuous neurofeedback or feedback delivered as soon as data is acquired, generally every TR (∼1-2 seconds), has the advantage of delivering somewhat immediate (relative to the 2-6 second hemodynamic delay) feedback, which may be important to aid in efficient learning. For continuous rtfMRI neurofeedback the hemodynamic delay must be accounted for in the feedback display or explained to subjects. A disadvantage of continuous neurofeedback using rtfMRI is that single measurements are often noisy, leading to the potential for inaccurate feedback if delivered on a TR-by-TR basis.

With continuous neurofeedback, subjects are asked to simultaneously attend to the task and the feedback signal, which increases cognitive load for the subject, and may disrupt optimal learning (van Merrienboer and Sweller, 2005). Other brain systems will be engaged by feedback that may be unrelated or even counterproductive to the task or training. For example, in a study by Greer and colleagues (Greer et al., 2014), subjects were better able to use neurofeedback to increase nucleus accumbens activity by visualizing exciting events than to decrease activity by visualizing boring events. Subjects may have been less capable of decreasing nucleus accumbens activity with neurofeedback because the reward response to successful self-regulation of brain activity conflicted with the concurrent task of visualizing neutral or non-arousing events. In such cases, it could be advantageous to delay neurofeedback.

Additional brain regions appear to be recruited related to neurofeedback itself which have not yet been well-characterized. A study by Haller and colleagues (Haller et al., 2013) reported increased functional connectivity during neurofeedback between an auditory target region and low level visual, insula, and working memory networks that was not found during a transfer phase with no feedback. Another study found activations in a broad frontoparietal and insula network as well as a broadly distributed negative network when subjects controlled neurofeedback as compared to viewing the feedback with no control (Papageorgiou et al., 2013). Control over feedback was further associated with improved whole-brain task signal-to-noise and increased pattern classification accuracy. Whether additional brain regions activated during neurofeedback are related to attention, control of feedback, or other factors such as learning and memory, has yet to be determined, including how these brain regions interact to produce neurofeedback-related brain change. This work highlights the need for studies which investigate brain changes during continuous neurofeedback to consider the confounding effects of the neurofeedback (and learning) itself. Finally, there is evidence that continuous feedback may interfere with the consolidation of a learned response. Animal and human studies of operant learning have shown improved learning when subjects are provided with a period of delay after reward for “post-reinforcement synchronization” (Sherlin et al., 2011).

Intermittent neurofeedback or neurofeedback delivered after data acquisition over several TRs, generally between 8-60 seconds, allows for averaging of the feedback signal to improve SNR and accommodate the hemodynamic delay, and also minimizes cognitive load and other potential confounds by separating task strategy from the evaluation of feedback. Johnson and colleagues found that subjects were better able to manipulate activity in the premotor cortex when imagining movements with intermittent neurofeedback (at the end of each 20 second block) as compared to continuous neurofeedback (Johnson et al., 2012). However, the optimal delivery of neurofeedback likely depends on the specific application. While intermittent neurofeedback might be more effective by reducing cognitive load in studies in which individuals are provided with specific practice strategies, continuous neurofeedback might be useful to train individuals to fine-tune cognitive strategies related to specific patterns of brain activity.

Finally, it is important for researchers to consider human-computer interface design principles, especially as they relate to the display of the neurofeedback signal in order to aid in effective learning. This is a critical consideration as poor human-computer interface design alone could lead to failed trials, or in some cases, adverse consequences such as increased frustration, confusion, and/or fatigue. No studies have been conducted to evaluate or optimize rtfMRI neurofeedback interface design. A review of human-computer interface design is beyond the scope of this paper (see Brown, 1998 for a general overview of this topic; Wickens et al., 2004); however, one area where human-computer interface design may be helpful is choosing the optimal modality for delivering neurofeedback. Most rtfMRI neurofeedback paradigms, to date, have used visual feedback, but some subjects and populations may benefit from auditory, haptic, virtual reality/immersion, or some combination of these modalities for neurofeedback. In one application, an EEG-based neurofeedback signal was displayed on a participant’s head using 3-dimensional cameras and a mirror to create a more realistic individualized display of brain function in real time (Mercier-Ganady et al., 2014). Future studies might query user experience (e.g., by questionnaires) to evaluate the functionality of the neurofeedback and the ease to which the user could learn to control the feedback signal, and compare this with a measure of accuracy with which the user could control the feedback, i.e., performance.

### Guideline 4: There is an appropriate test of training success

It is also important to establish guidelines for how best to assess rtfMRI training success. To date, there have been two common approaches: subjects show improved (1) control of brain activity while receiving neurofeedback and/or (2) control of brain activity without neurofeedback (i.e., by comparing brain regulation without neurofeedback before and after training with feedback). A few studies have tested for transfer from training runs in which cognitive strategies were provided to control the feedback, to transfer runs in which cognitive strategies were used to control brain activity without any fMRI information. Young and colleagues reported that after several runs of neurofeedback training to increase activity in the left amygdala, subjects were able to significantly increase activity in the left amygdala using only cognitive strategies without neurofeedback (Young et al., 2014). In another recent study, Robineau and colleagues (Robineau et al., 2014) trained subjects to control interhemispheric visual cortex balance over three hour-long neurofeedback training sessions comprised of both feedback and transfer runs. They found that participants who learned to control the feedback signal were able to maintain control during transfer runs with no neurofeedback. In other studies, subjects had success self-regulating brain activity with the aid of neurofeedback, but they were not able to self-regulate activity when no longer receiving neurofeedback (Berman et al., 2013; Caria et al., 2007; Greer et al., 2014; Hamilton et al., 2011). When creating an operational definition for training success, it will be important to consider: the expected timescale of training effects, expected pattern of change (e.g., linear or non-linear, monotonic or non-monotonic), and how best to account for individual differences. For example, training effects may be observed immediately or following some delay depending on the type and nature of learning impacting or impacted by training. Successful training may occur via gradual, incremental improvement characterized by a linear, monotonic function or trial-and-error testing characterized by a non-linear, non-monotonic function. It is also unclear whether the experimenter should fix the training interval or allow for adaptive training based on individual differences in optimal learning strategies and performance. Finally, the experimenter will need to design the study to adequately capture potential brain changes related to training over time, which could include reduced activity in the target ROI(s), change in the extent of activation within the ROI(s), and/or recruitment of different neural systems to support improved performance. The experimenter could consider including a resting state fMRI scan before and after training as one strategy for capturing complex brain changes over time (Hampson et al., 2011; Scheinost et al., 2013).

In order to determine whether rtfMRI neurofeedback training-induced changes are *clinically* significant, it will be important to introduce a metric of change such as the reliable change index (Jacobson and Truax, 1991) that accounts for measurement error and determines how functioning compares to a normative population. This is often used in clinical and neuropsychological assessments/studies where clinical significance is a question. Neuropsychological and clinical measures used for this type of an assessment typically have established psychometric properties (normative data, test-retest reliability, etc.), which make calculating reliable change indices possible. As the Human Connectome Project and other large neuroimaging databases are developed and data is shared, normative databases could be developed for fMRI data in order to calculate a reliable change index for neuroimaging data.

### Guideline 5: rtfMRI neurofeedback leads to behavioral change

The behavioral effects of rtfMRI training may be manifest in improvement on the task used during rtfMRI neurofeedback training, improvement on related tasks or on the same task in other contexts, or improvement that generalizes to real-world outcomes (for review, see Ruiz et al., 2014).

In some studies, subjects have been trained to self-regulate brain activity by manipulating the neurofeedback signal and a behavioral response to some other, often concurrent stimulus. For example, deCharms and colleagues reported that training self-regulation of activity in the dorsal anterior cingulate, a brain region implicated in pain perception and regulation, led to a corresponding change in the perception of pain caused by a noxious thermal stimulus as well as in spontaneous pain perception in patients with chronic pain (deCharms et al., 2005).

Another approach is to assess behavioral change before and after rtfMRI feedback training. For example, Zhang and colleagues trained subjects to increase activation in the left dorsolateral prefrontal cortex, a brain region involved in working memory, and reported improvements on digit span and letter memory tasks across training, indicating improved verbal working memory with rtfMRI neurofeedback training (Zhang et al., 2013). However, another study by Lawrence and colleagues found that although rtfMRI feedback could be used to train subjects to increase activity in the right anterior insula, training did not lead to changes in skin conductance response or subjective valence ratings across pre- and post-training affective probes (Lawrence et al., 2013). Because so few evidence-based guidelines exist for rtfMRI studies, it is difficult to determine whether a lack of behavioral change is related to the specific brain-behavior relationship tested, or to methodology, such as only one rtfMRI training session in that study.

There has been limited work in non-clinical populations targeting emotional brain regions and function. Rota and colleagues have shown improved detection of emotional tone with rtfMRI training from the right inferior frontal gyrus in healthy participants (Rota et al., 2009). On the other hand, another study (Johnston et al., 2011) found that rtfMRI training from brain regions involved in positive emotions failed to improve mood ratings in healthy participants. Although methodology must be considered, the authors suggest that rtfMRI training of emotional control to enhance mood may be most effective in individuals with abnormal emotional control (as in their prior study in depression Linden et al., 2012), and less effective in individuals who are capable of normal mood regulation (but see Allen et al., 2001 for a study demonstrating improved mood in health controls using EEG neurofeedback). A recent study of individuals with major depressive disorder (Young et al., 2014) found that rtfMRI neurofeedback training to increase activity in the amygdala during positive autobiographical memory recall led to improved self-reported mood post-scan compared to controls.

There is limited evidence of behavioral change from rtfMRI that has generalized to other tasks or real-world outcomes. Prior studies in clinical populations have shown decreased pain ratings in individuals with chronic pain (deCharms et al., 2005), decreased symptoms in individuals with tinnitus (Haller et al., 2010), decreased craving ratings and physiological response to smoking cues in nicotine-dependent individuals (Canterberry et al., 2013; Hanlon et al., 2013), decreased mood symptoms in people with depression (Linden et al., 2012), increased motor speed and clinical ratings of motor symptoms in individuals with Parkinson’s disease (Subramanian et al., 2011), and decreased contamination anxiety in people with sub-clinical anxiety (Scheinost et al., 2013). However, with one exception (Scheinost et al., 2013), these studies measured behavior at the time of the rtfMRI study, did not test retention, and, with few exceptions (Scheinost et al., 2013; Subramanian et al., 2011) the dependent measure of behavioral change with rtfMRI was assessed with self-report measures that may reflect non-specific training effects.

Moving forward, it will be important for experimenters to show a causal link between the brain region(s) trained and the behavior targeted for modification in order to establish that specific brain-behavior changes account for any observed clinical changes and are not simply epiphenomena. It is a challenge to demonstrate cause-and-effect beyond correlation between rtfMRI neurofeedback training and behavioral change. Related to this, researchers should consider the specificity of the hypothesized relationship between the brain activation pattern(s) trained and behavioral changes. In the study by Shibata and colleagues described above (Shibata et al., 2011), for example, causality was established between rtfMRI neurofeedback training from early visual cortex corresponding to a specific visual orientation and improved accuracy in identifying the target orientation as compared to untrained orientations. In such studies the specific brain activation pattern related to behavior must first be determined, for example using multivariate pattern analysis as in Shibata et al., and as described above (LaConte, 2011). In another recent study, Zhao and colleagues (Zhao et al., 2013) used dynamic causal modeling with rtfMRI to measure causal interactions between the dorsal premotor cortex target region and other motor regions during a motor imagery task with neurofeedback. The experimental group showed increased interactions from the target brain region to the other regions across four training sessions as compared to a sham control group. The experimental group also showed improved performance on the motor imagery task. rtfMRI methods need to be further refined to establish causality not only between neurofeedback training and changes in behavior, but also between changes in the specific neurofeedback signal and other brain changes.

### Guideline 6: An appropriate rtfMRI neurofeedback-based clinical trial design is in place

Despite important early work suggesting it is possible to use rtfMRI as a non-invasive brain-based clinical tool, to our knowledge there has been only one RCT, recently completed, that has investigated the efficacy of rtfMRI neurofeedback to effect meaningful clinical change (CH, personal communication). Several methodological considerations for the design of rtfMRI studies and clinical trials remain open questions that likely depend on the specific application.

To demonstrate behavioral change that is directly related to rtfMRI feedback training and establish causality, studies must implement important control conditions. Most studies that have included a control condition have either used false feedback or no feedback. False feedback can involve providing subjects with arbitrary feedback not related to brain function, actual neurofeedback from a brain region or network theoretically unrelated to the experimental variables of interest either within-group (e.g., Garrison et al., 2013b) or between groups (e.g., Lawrence et al., 2013; Young et al., 2014), or yoked neurofeedback from a matched subject (e.g., Hampson et al., 2012). In a recent study, false feedback based on a recording of EEG neurofeedback was found to engage a broad network of frontal, parietal and cingulate regions involved in cognitive control (Ninaus et al., 2013). In another study, fixed randomized feedback not based on actual fMRI signal was compared to no feedback as a control for training self-regulation of activity in the premotor cortex (Johnson et al., 2012). False feedback again produced a widespread pattern of activation involving frontal, temporal, and parietal regions, which was distinct from the more localized activation associated with real neurofeedback. In addition, subjects reported more frustration with the task in the false feedback group as compared to the no feedback or real neurofeedback groups. Based on these findings, the authors reasoned that the negative impact of false feedback runs made it a less suitable control group than no feedback. However, the drawbacks to providing no feedback are first that it is unlikely to be as engaging as a feedback task, and second, that it does not control for the perception of success that subjects experience when they do well controlling their brain patterns. These differences between a neurofeedback group and a no feedback control group can lead to false positives related to unmatched motivation and placebo effects. Another option is to deliver control feedback using more cost-effective methods such as autonomic biofeedback (deCharms et al., 2005).

Other important design considerations for rtfMRI clinical trials include the optimal number of rtfMRI sessions, number of neurofeedback runs per session, appropriate timing between sessions if multiple sessions are used, and the combination of rtfMRI training with behavioral or other interventions. There is little data available to guide optimization of these parameters for clinical trials. A recent study addressed a number of these issues by tracking change across three rtfMRI sessions in which nicotine-dependent individuals were trained to reduce activation in the anterior cingulate cortex and reduce smoking cue-related craving (Canterberry et al., 2013). Of 15 enrolled smokers, sixty percent completed three 1-hour rtfMRI sessions, 1-2 weeks apart. Within each rtfMRI session, subjects completed three 10-minute feedback runs. Reduced anterior cingulate cortex activity and reduced self-reported craving were evident at the first rtfMRI session and consistent across sessions and runs. This reduction in cue-induced craving with rtfMRI neurofeedback was significant at the third session, indicating that at least two feedback sessions were necessary to see any effect of neurofeedback, and more than two sessions may be needed to observe clinical improvement.

Finally, it will be critical to compare the effects of rtfMRI-based neurofeedback to existing therapies or biofeedback using more cost-effective neuroimaging tools such as EEG in order to demonstrate the value added by rtfMRI-based neurofeedback above other treatment options.

### Guideline 7: Sharing resources and using common standards

In a domain where reproducibility has been a non-trivial goal, there is a need for consensus on common standards and sharing of data, paradigms, software, and analytic tools. This will provide an additional benefit of lowering the barrier to entry for researchers to use rtfMRI, which will also help generate new research questions and the development of novel algorithms, solutions, and tools to advance the field.

One area of importance is the creation of an open rtfMRI communication protocol. Although Digital Imaging and Communications in Medicine (DICOM) is a standard for communicating imaging data to Picture Archiving and Communication System (PACS) and other systems, many scanners do not have the capability to reconstruct and send DICOM images as they are acquired. As such there is no common standard across scanners to communicate rtfMRI data from the scanner to an analysis or presentation computer. In the absence of such standards, several existing software packages rely on monitoring the file system to detect reconstructed images. While simpler to setup, this can introduce unnecessary delays and limit the possibilities of neurofeedback paradigms. In scanners manufactured by Siemens (www.siemens.com) and Philips (www.philips.com), users can transmit data over the network. In recent work published openly on Github (http://github.com/gablab/murfi2), a formal specification for rtfMRI communication has been developed to describe a set of information necessary to be transmitted from the scanner to an analysis or presentation computer. Software developers can use this standard to create both new sequences on scanners as well as new analysis platforms that communicate with these sequences. However, if manufacturers were able to send DICOM images in real time, it would benefit from an established protocol.

Along with available standards, a repository for or notification of the availability of such resources is needed. A common neuroinformatics portal for rtfMRI coupled with question-answer sites (e.g., http://neurostars.org) and code repositories (e.g., GitHub, http://github.com) can significantly simplify the dissemination of information and allow for community discussion of approaches and issues. With the increased focus on data sharing and reproducibility of imaging studies, it is critical to utilize such resources to increase sharing of rtfMRI data, experimental paradigms, and software.

## Potential impact of real time fMRI for clinical neurotherapeutics

A primary goal of current research using rtfMRI neurofeedback is to aid in the development of safe, effective and personalized therapies for many brain related disorders including: pain, addiction, phobia, anxiety, and depression. rtfMRI neurofeedback has direct clinical application as a standalone treatment or as an augmentation strategy for interventions that work by training volitional control of brain activity. As described, rtfMRI neurofeedback has been used to train individuals to self-regulate brain activation patterns related to basic and clinical processes, and RCTs are in progress to corroborate potential clinical outcomes. rtfMRI may be especially effective as targeted neurofeedback in conjunction with behavioral interventions based on brain-behavior relationships. Additionally, and on the other end of the central nervous system disease spectrum, there is the potential application of rtfMRI neurofeedback to enhance performance (UK Mindfulness-Based Teacher Trainer Network, 2011), learning (Yoo et al., 2012), perception (Scharnowski et al., 2012), or to promote wellness optimization.

In clinical neurotherapeutics, improvements in our understanding of the neural underpinnings of psychiatric disorders have yielded potential neural targets for rtfMRI interventions. For example, dysfunction of the subgenual cingulate has been implicated in refractory depression, and deep brain stimulation targeted to that brain region has shown preliminary efficacy for the treatment of depression (Mayberg et al., 2005). As testing the efficacy of deep brain stimulation is invasive, it is possible that rtfMRI may be used for targeted neurofeedback to test this hypothesis prior to more invasive procedures. More generally, rtfMRI has the potential to test the robustness of neurobiological hypotheses prior to more invasive procedures.

Looking forward, there is a clear need for cost-effective therapies. At this time, rtfMRI neurofeedback is costly. However, rtfMRI research can inform the development of more cost-effective and scalable clinical tools, such as EEG or functional near-infrared spectroscopy (fNIRS). A translational step is to test multimodal approaches such as simultaneous rtfMRI-EEG. In a recent study using this approach, participants were able to simultaneously increase BOLD signal in the amygdala and frontal high-beta EEG asymmetry when provided both modes of neurofeedback during a positive emotion induction task (Zotev et al., 2014), a technique which may help improve mood regulation in individuals with major depressive disorder (Young et al., 2014). The combined rtfMRI-EEG approach may provide more efficient training based on improved temporal and spatial resolution compared to either modality alone. The concurrent use of these modalities may also help to translate rtfMRI neurofeedback into more feasible EEG or other imaging applications (e.g., fNIRS) by characterizing the EEG (or fNIRS) signal correlated to the fMRI signal of interest to be used for neurofeedback.

## Real time fMRI in clinical neuroimaging beyond neurotherapeutics

Resolution of the issues outlined here will contribute to the development and use of rtfMRI as a clinical neuroimaging tool beyond the direct therapeutic applications of rtfMRI neurofeedback that are the focus of current rtfMRI research and likewise of this paper. In addition to the use of rtfMRI as a tool for treatment via neurofeedback, rtfMRI has potential utility in exploring the nature of the pathological condition, and in clinical diagnosis, monitoring disease course, and tracking therapeutic response (including the effects of neurofeedback). rtfMRI offers a significant new opportunity for understanding and addressing these broad clinical problems.

At the basic or translational level, rtfMRI can be used to clarify brain-behavior relationships critical to the understanding and treatment of brain disorders. In particular, rtfMRI can be used to improve our understanding of how cognitive processes are represented in the brain and how cognition is related to behavior in real time. For example, subjective information can help to elucidate cognition (or disordered cognition), yet traditional self-report measures have limitations. rtfMRI can be used to relate subjective experience to objective neuroimaging data to gain a more complete understanding of these brain-behavior relationships, including in individual subjects. A recent study by Garrison and colleagues (Garrison et al., 2013a; Garrison et al., 2013b) used rtfMRI in this way to link the subjective experience of focused attention to brain activity in the default mode network in experienced meditators. Short fMRI task runs and immediate self-report were paired with offline feedback (shown after self-report), real time feedback, or volitional manipulation of the feedback stimulus. Meditators reported that their subjective experience corresponded with feedback, and showed a significant percent signal change in the target brain region upon volitional manipulation, confirming their reports. This approach obviates the problem of reverse inference whereby cognitive processes are inferred from brain activity (Poldrack, 2006), and reduces the opacity of cognitive strategy in fMRI studies. However, as meditators have been shown to be more accurate at introspection than non-meditators (Fox et al., 2012), the accuracy of self-report must be considered when determining these brain-behavior relationships in other groups, including clinical populations. Nevertheless, rtfMRI may be used in this way to help determine the specific brain activation pattern(s) related to cognitive processes or behaviors of interest. This approach may aid in rtfMRI neurofeedback training and provide insight into the mechanisms of treatment in RCTs using rtfMRI neurofeedback. More generally, this use of rtfMRI may further our understanding of cognitive processes including those relevant to clinical applications.

As a tool in clinical neuroimaging, rtfMRI neurofeedback has the potential to be used to inform clinical diagnosis, track the natural history of disease, track treatment progress, and provide more specific and effective treatment. The use of rtfMRI neurofeedback could lead to individualized brain-based treatment by clarifying the neural underpinnings of disordered behavior in an individual through *real time* testing. This could be accomplished: (1) by manipulating different cognitive processes that may be disrupted in a given disorder to observe the change in brain function in real time; (2) by presenting a cognitive task when different patterns of brain function are observed in real time; and (3) by altering brain function in different regions with reversible brain stimulation tools such as transcranial magnetic stimulation (TMS) to determine the effects on brain function, cognition, and clinical symptomatology. rtfMRI neurofeedback may then be used to modify these disordered individual patterns of brain activation toward a more normative state, potentially leading to more healthy, adaptive brain function. rtfMRI neurofeedback may be calibrated to the individual’s current state, for example, to enhance learning as an individual improves across an intervention. Likewise, interventions may be tailored to the specific strategies found to be useful for the individual in rtfMRI neurofeedback studies (Marchand, 2013). Further, behavioral interventions may be augmented by targeted neurofeedback of a brain activation pattern or cognitive process of interest. Finally, rtfMRI may be used to find predictors of not only clinical outcomes, but also of responsiveness to rtfMRI neurofeedback training.

## Conclusions

This paper recommends issues for consideration in studies of rtfMRI neurofeedback for clinical therapeutics. Research contributing to evidence-based guidelines is sorely needed for clinical trials of rtfMRI neurofeedback. As outlined, researchers must establish that the rtfMRI signal is accurate and reliable, rtfMRI neurofeedback leads to learning, there is an appropriate test of training success, rtfMRI neurofeedback leads to clinically meaningful changes in cognition and behavior, an appropriate clinical trial design is in place, and rtfMRI resource sharing protocols and tools are established to allow for efficient advancement toward the urgent clinical goals discussed in this paper. Important beginning work in these areas has been conducted, contributing to an overall promising outlook for the application of rtfMRI neurofeedback to develop novel, safe, and effective treatments for brain disorders. The ultimate goal is for this tool to assist clinicians and patients in designing personalized assessment and intervention approaches that may enhance resilience in at-risk populations by correcting known maladaptive patterns of brain function in advance of developing a disorder, accelerating adaptive compensatory neuroplastic changes in those with brain disorders, and/or directly targeting the disrupted brain region or system underlying brain disorders in order to restore healthy brain-behavior function. Overall, rtfMRI offers the opportunity to further our understanding of how the brain works and pushes the limits of our potential for self-directed healing and change.

## Acknowledgments

The ideas for this paper were developed at a workshop in July 2013 supported by the Radcliffe Institute for Advanced Study. Support was also provided in part by the National Institutes of Health (R21/33DA026104 (HCB, AK); K23DA032612 (LES); K01 DA034093 (JMG), K24DA030443 (AEE); K24DA029262 (SM), R01 AT007922-01 (JAB), R03 DA029163-01A1 (JAB), K12 DA00167 (JAB), P50 DA09241 (JAB), R01 AA02152901A1 (SML), R01 MH095789 (MH), 5R21DA026085 (MG), Poitras Center for Affective Disorders Research (SWG), the Charles A. King Trust (LES), the Norman E. Zinberg Fellowship in Addiction Psychiatry at Harvard Medical School (LES, JMG), the McGovern Institute Neurotechnology Program (SG, JDG, AEE), and private funds to the MGH Department of Psychiatry (AEE). Research presented in this paper was also carried out in part at the Athinoula A. Martinos Center for Biomedical Imaging at the Massachusetts General Hospital, using resources provided by the Center for Functional Neuroimaging Technologies, P41RR14075, a P41 Regional Resource supported by the Biomedical Technology Program of the NIH National Center for Research Resources (NCRR).

## Footnotes

1 Most rtfMRI systems use blood oxygen-dependent level or BOLD contrast, which is an indirect measure of brain function with known spatial and temporal resolution limitations.

2 Real time, in the context of real time fMRI, refers to the ability to capture a brain signal every 1-2 seconds with the limitation that the BOLD response takes 2-6 seconds to rise to peak LaConte, S.M., 2011. Decoding fMRI brain states in real-time. Neuroimage 56, 440-454, Logothetis, N.K., Pauls, J., Augath, M., Trinath, T., Oeltermann, A., 2001. Neurophysiological investigation of the basis of the fMRI signal. Nature 412, 150-157.

